# Serotonin drives choice-independent reconfiguration of distributed neural activity

**DOI:** 10.1101/2025.08.01.668048

**Authors:** Guido T. Meijer, Joana A. Catarino, Laura Freitas-Silva, Inês Laranjeira, International Brain Laboratory, Zachary F. Mainen

## Abstract

Serotonin (5-HT) is a central neuromodulator which is implicated in, amongst other functions, cognitive flexibility. 5-HT is released from the dorsal raphe nucleus (DRN) throughout nearly the entire forebrain. Little is known, however, about how serotonin affects downstream populations of neurons and how this modulation might support its cognitive functions. Here, we optogenetically stimulated serotonergic neurons in the DRN while recording large parts of the brain with Neuropixels during quiet wakefulness and performance of a perceptual decision-making task. During quiet wakefulness, 5-HT stimulation induced a rapid switch in internal state, as indicated by dilation of the pupil, suppression of hippocampal sharp wave ripples, and increased exploratory behaviors, such as whisking. To elucidate the brain-wide effect of serotonin release we performed acute Neuropixel recordings in seven target locations, a total of 7,478 neurons were recorded across 17 mice. We found that 5-HT stimulation significantly modulated neural activity in all the recorded brain regions, both during quiet wakefulness and task performance. During task performance, however, we observed no change in behavior when stimulating 5-HT. We found that the 5-HT modulation of high-dimensional neural dynamics is confined to a subspace which is orthogonal relative to the choice axis. These observations describe a possible mechanism for the induction of state-dependent stimulus representations, suggesting a neural basis for neuromodulatory effects on brain-wide circuits to flexible decision-making.

## Introduction

Serotonin (5-hydroxytriptamine; 5-HT) is a central neuromodulator which is implicated, amongst other functions, in the regulation of cognitive flexibility^1–4^. 5-HT is released from the brainstem dorsal raphe nucleus (DRN) which projects throughout the entire brain^5,6^. The cognitive function of serotonin signalling in the brain is a topic of active debate. Serotonin has been suggested to code for reward/punishment^7–11^, surprise^1,3,12^, and patience/persistence^13,14^. The heterogeneity in the results of studies on serotonin make it difficult to pin down a single function for serotonergic signalling in the brain, this is exacerbated by the observation that the effects of serotonin stimulation are context specific^15,16^. Also at the neuronal level, the effects of serotonergic modulation vary widely in the literature ranging from inhibitory^17–19^ to excitatory^8^. Taken together, these studies do not paint a cohesive picture of what serotonin signals and how this signal modulates downstream target regions.

To shed light on this, we performed Neuropixel recordings in seven insertion targets while optogenetically stimulating 5-HT neurons in the DRN during quiet wakefulness and task performance. Given serotonin’s supposed role as a surprise signal we hypothesized that it would modulate how animals incorporate prior knowledge into current sensory evidence, and how fast their prior would update given changes in the environment. Contrary to our hypothesis, we did not find any effect of 5-HT stimulation on these behaviors. Even though a pulse of serotonin elicited strong behavioral changes during quiet wakefulness and widespread bidirectional modulation of neural dynamics across the brain.

How can strong neuronal effects be observed without a corresponding behavioral impact? We found that serotonergic modulation of neural dynamics was confined to a subspace which was orthogonal to the choice axis. This allows for the coding of modulatory changes during task performance which do not interfere with the execution of the task at hand. Such a coding scheme can potentially aid flexible behavior, whereby sudden changes in the environment require rapid updating of neural representations which drive subsequent behavioral change.

### Serotonin stimulation elicits a shift in internal state during quiet wakefulness

We selectively targeted serotonergic neurons in the DRN by injecting a Cre-dependent virus, coding for channelrhodopsin (ChR2), into the DRN of SERT-cre mice. To stimulate 5-HT neurons, an optical fiber was implanted over the DRN through the cerebellum (Fig. 1a). Expression was confirmed to be confined to the DRN in histology (Fig. 1b). The relative fluorescence in the DRN, compared to a control area in the same slice, was only elevated in SERT-cre mice (n=11) and not in wild-type controls (n=6; Fig. 1c; see Supp. Fxig. 1 for histology of all mice). Mice were head-fixed in a rig set up such that mice could perform the steering wheel task during acute dual Neuropixel insertions (Fig. 1d). A recording session started with allowing the probes to settle in the brain, after which the mouse performed the behavioral task, followed by six minutes of spontaneous activity, and concluded by passive stimulation of 5-HT during quiet wakefulness (Fig. 1e).

**Figure 1.**
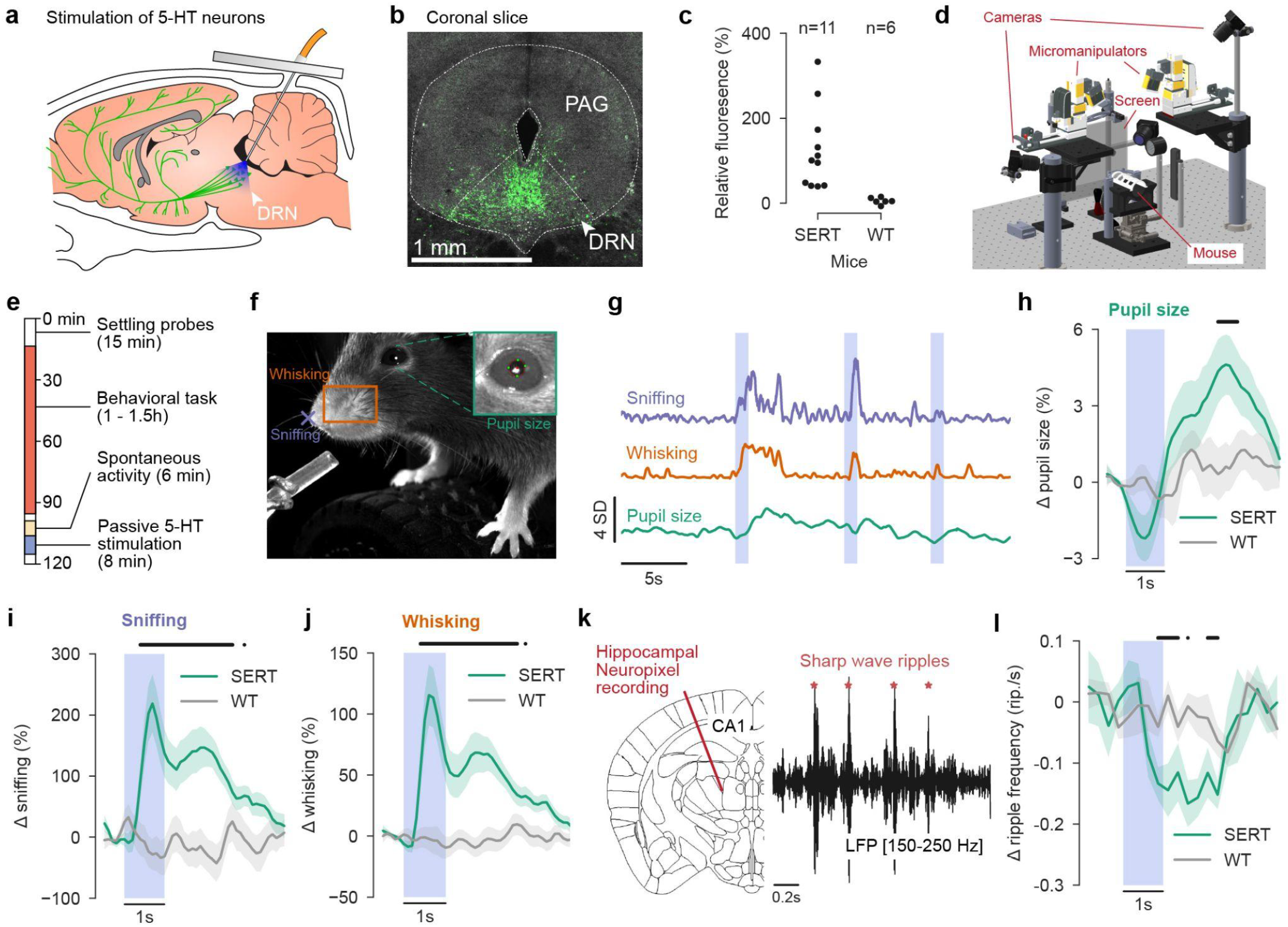
Serotonin stimulation induces a switch in internal state during quiet wakefulness. **(a)** A cartoon depiction of 5-HT neurons in the dorsal raphe nucleus (DRN) which project throughout the brain. An optical fiber is implanted at an angle, through the cerebellum. **(b)** Channelrhodopsin was targeted to serotonergic neurons in the DRN using a viral injection, localized expression was confirmed with histology: green is the co-expressed GFP marker. **(c)** The relative fluorescence in the DRN, compared to a control region in the same slide, for sert-cre (SERT) and wild-type control (WT) mice. **(d)** 3D render of the Neuropixel recording setup including two Sensapex manipulators and three cameras. **(e)** A session started with letting the inserted probes settle in the brain after which the behavioral task was run. After the task six minutes of spontaneous activity was recorded followed by the passive stimulation of 5-HT. **(f)** From the video several physiological markers were extracted. Pupil size was defined as the circle which intersects four tracked DLC points around the pupil, whisking as the movement energy in an ROI around the whisker pad, and sniffing as the frame-to-frame movement of a tracked DLC point on the tip of the nose. **(g)** Example snippet of the physiological markers from (a), blue vertical bars indicate 1s 5-HT stimulation during quiet wakefulness. **(h)** The change in pupil size, relative to baseline, conditioned on the start of passive 5-HT stimulation (blue vertical bar), for sert-cre (SERT) and wild-type control (WT) mice. Black horizontal bar indicates a significant difference between SERT and WT mice (*p* < 0.05, independent samples t-test). Green and grey thick lines are the means over animals and the shaded region indicates the s.e.m. over animals. **(i)** Same as (h) but for whisking. **(j)** Same as (h) but for sniffing. **(k)** Detection of hippocampal sharp wave ripples from CA1 local-field potential. **(l)** Same as (c) but for hippocampal sharp wave ripple frequency.

We first focussed on the effects of serotonin stimulation during quiet wakefulness. Specifically, we replicated the previous finding that serotonin stimulation results in an increase in pupil size^20^. Pupil size was tracked from the video feed (Fig. 1f,g) and centered at passive 5-HT stimulation bouts during quiet wakefulness. A significant increase in pupil size was observed in SERT-cre mice compared to WT controls ∼2 seconds after 5-HT stimulation offset (Fig. 1h). Besides an increase in pupil size, 5-HT stimulation also elicited exploratory behaviors (i.e. sniffing and whisking) at short latency (Fig. 2i,j). These observations together suggest that the animal shifts from an “offline” state to a more “online” state after a transient pulse of serotonin. To further corroborate this claim, we investigated the effect of 5-HT stimulation on hippocampal sharp wave ripples (Fig. 1k). Sharp wave ripples occur during sleep, but also when the animal is awake but in a quiescent state, associated with low locomotion and small pupil size^21–23^. Indeed we found that 5-HT stimulation resulted in a significant reduction of hippocampal sharp wave ripples (Fig. 1l), suggesting a shift from an offline to an online state.

**Figure 2.**
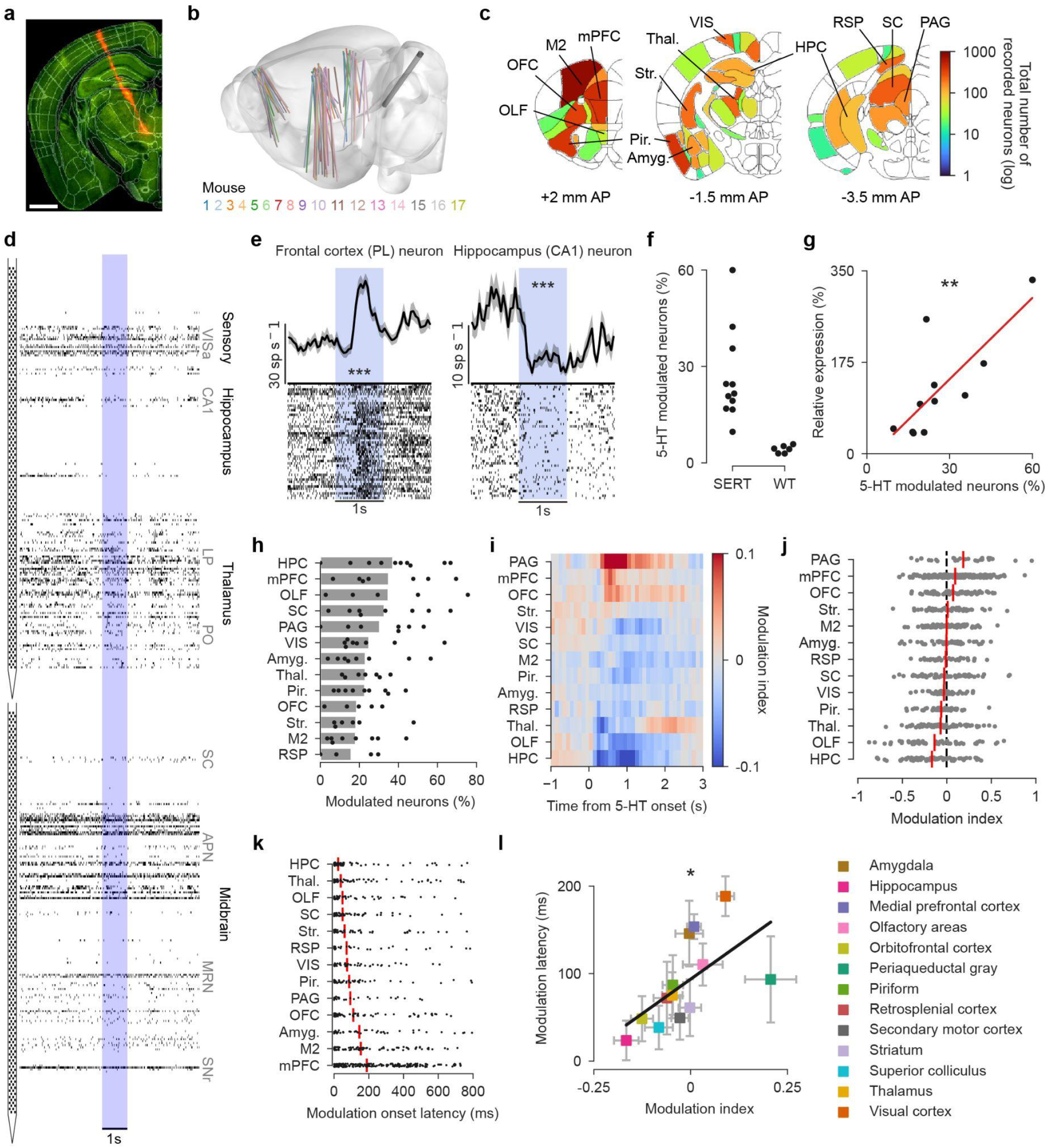
Serotonin stimulation bidirectionally modulates neural dynamics across the brain during quiet wakefulness. **(a)** Neuropixels were coated with Di-I (red trace) to allow post-hoc reconstruction of the probe tract. White bar indicates 1 mm. **(b)** All Neuropixel insertions, tracked in Allen atlas space, and plotted in a standardized brain. The color indicates from which animal each insertion is from. **(c)** Total number of recorded neurons per brain region. OLF: olfactory areas, OFC: orbitofrontal cortex, M2: supplementary motor cortex, mPFC: medial prefrontal cortex, PIR: piriform cortex, Str.: striatum, Amyg.: amygdala, Thal.: thalamus, VIS: visual cortex, HPC: hippocampus, RSP: retrosplenial cortex, SC: superior colliculus, PAG: periaqueductal grey. **(d)** Snippet from an example dual-Neuropixel recording, each tick is a spike from a clustered unit that passed all QC criteria. The blue vertical bars indicate when the optogenetic stimulation of 5-HT neurons was on. On the right are the recorded brain regions, at their respective depths along the probes. VISa: anterior visual cortex, LP: lateral posterior nucleus of the thalamus, PO: posterior complex of the thalamus, SC: superior colliculus, APN: anterior pretectal nucleus, MRN: midbrain reticular nucleus, SNr: substantia nigra, reticular part. **(e)** Two typical examples of 5-HT modulated neurons. The black line shows the mean firing rate over trials (50 ms bins with 25 ms Gaussian smoothing window), with gray shading indicating the standard error of the mean. The bottom part of the plot shows the spike raster in which each row is a stimulation trial with ticks showing individual spikes. The blue rectangle indicates the time at which the optogenetic stimulation was on. ***: ZETA test *p* < 0.001. **(f)** Percentage of significantly modulated neurons by 5-HT stimulation per recorded animal (each dot is a mouse), plotted separately for sert-cre (SERT) and wild-type control (WT) animals. **(g)** The percentage of 5-HT modulated neurons was correlated with the amount of expression, as determined from histology. ** *p*=0.005, r=0.77, Pearson correlation. **(h)** Fraction of significant 5-HT modulated neurons per brain region (*p* < 0.05, ZETA test). Black dots indicate individual mice (only sert-cre animals are plotted), grey bars show the mean over animals. **(i)** The modulation index, defined as the ratio of firing rate change relative to baseline (last time bin before 5-HT stimulation onset), calculated for 100 ms non-overlapping time bins. Modulation index is computed for all significantly modulated neurons and averaged per brain region. Red colors indicate an increase in neural activity and blue colors a decrease relative to baseline. **(j)** The modulation index plotted per neuron (grey dots), defined as the ratio between a baseline [-0.5s - 0s] and a 5-HT window [0.3s - 0.8s]. Red vertical lines are the mean over all neurons per brain region. **(k)** The temporal onset of 5-HT modulation, defined by the ZETA test, for each neuron (black dots) averaged per region (red vertical lines). **(l)** Regions were suppressed at short time latencies and enhanced at longer latencies. Each colored square is a brain region with the mean modulation index of all neurons in that region plotted on the x-axis and the s.e.m. plotted as grey whiskers, same for the modulation latency on the y-axis. * *p*=0.028, r=0.61, Pearson correlation between mean latency and modulation index per region.

### Serotonin modulates neural dynamics across the brain during quiet wakefulness

To investigate how 5-HT modulates neural activity in downstream target regions, we performed acute Neuropixel recordings during optogenetic serotonin stimulation. The positioning of the probes in the brain was post-hoc confirmed with histology by tracing the fluorescent tract left by the probe, which was coated in Di-I (Fig. 2a). Typically two probes were inserted at the same time to record from hundreds of neurons across many brain regions simultaneously. Preprocessing and spike sorting was performed using the International Brain Laboratory (IBL) pipeline^24^, after which neurons were localized to brain regions by aligning the electrophysiological signatures along the probe with histology. A total of 7,478 neurons were recorded in 13 brain regions (n=86 probe insertions during 57 recording sessions in 17 mice; Fig. 2b-d).

Single neurons showed diverse patterns of 5-HT modulation; some showed an increase in activity whereas others were suppressed (Fig. 2e; see Supp. Fig. 2 for activity patterns of all neurons). In SERT-cre animals, anywhere between 10 and 60% of neurons were significantly modulated while in wild-type controls the percentage of neurons was around chance level (5%; Fig. 2f). The percentage of 5-HT modulated neurons significantly correlated with the level of channelrhodopsin expression, as determined by histology (Fig. 2g). The amount of 5-HT modulated neurons varied by brain region, with the most modulated neurons in the medial prefrontal cortex and the hippocampus and the least in the retrosplenial cortex (Fig 2h). Despite this variability, all recorded brain regions showed a percentage of modulated neurons which was higher than the 5% chance level.

The sign of the modulation was similarly variable across brain regions. We computed a modulation index, which is positive when a neuron’s activity is elevated by 5-HT stimulation and negative when it’s suppressed, compared to baseline. Taking the mean modulation index over all neurons per brain region showed that the periaqueductal gray, medial prefrontal, and orbitofrontal cortex showed a transient increase in activity. Contrary to the thalamus, olfactory areas, and the hippocampus, which showed a suppression (Fig. 2i). The mean modulation index over neurons, however, does not reveal neuron-to-neuron differences in 5-HT modulation. Therefore, we plotted the modulation index per neuron for the time window of 300 - 800 ms after serotonin stimulation onset. This revealed a large variability over neurons, brain regions that were enhanced on average, also included many suppressed neurons and vice versa (Fig. 2j). Furthermore, brain regions which on average did not show an effect contained many enhanced and suppressed neurons whose effects cancelled each other out at the population level. In other words, these regions showed a balance between 5-HT induced excitation and inhibition.

Serotonin modulates neural activity through a wide range of serotonergic receptors, predominantly G-protein coupled receptors activating myriad intracellular signalling pathways, but also ionotropic receptors^25^. Besides high variance in the sign of the modulation, also the latency at which a pulse of 5-HT modulates neural activity in downstream regions is expected to be highly variable. Indeed, we found a lot of neuron-to-neuron variability in the onset latency of 5-HT modulation. The latencies, however, did show differences across regions (Fig. 2k). For example, hippocampus was modulated at short latency whereas medial prefrontal cortex showed longer latencies. There was a significant correlation between the mean onset latency of a region and its 5-HT modulation sign, such that inhibited brain regions showed short latencies while excited regions long latencies (Fig. 2l).

This begs the question whether the inhibition was mediated through rapid activation of inhibitory neurons. We separated narrow spiking neurons from wide spiking neurons using their spike width (Supp. Fig. 3a). Narrow spiking neurons are putative fast-spiking interneurons; we confirmed that the firing rates of narrow spiking neurons were significantly higher compared to wide spiking neurons (Supp. Fig. 3b). Serotonin stimulation modulated slightly more putative interneurons than other neurons (Supp. Fig. 3c). However, there was no significant difference between the modulation index of putative interneurons and the rest (Supp. Fig. 3d). In short, the rapid inhibition observed after serotonin stimulation was not due to excitation of fast-spiking interneurons.

Another possibility is that serotonin has different modulatory effects across the cortical layers. We split all our cortical recordings by layer, but observed no difference in the percentage of 5-HT modulated neurons per layer (Supp. Fig. 4a,b). The sign of the serotonergic modulation and the onset latency also did not show any layer dependent differences (Supp. Fig. 4c,d).

To conclude, serotonin significantly modulated neural dynamics in all recorded brain regions and did so in a bidirectional manner. Modulation onset latency correlated with modulation sign such that inhibition was fast whereas excitation was slow.

### Serotonin stimulation does not affect decision making

Having established that 5-HT stimulation induced a shift in internal state during quiet wakefulness and widespread modulation of neural activity across the brain, we expected that it would impact decision making. Since transient serotonin release is considered to be a surprise signal we hypothesized that 5-HT stimulation would affect the way mice incorporate prior knowledge with current sensory evidence. To test this, we trained mice in the steering wheel task in which stimuli were presented on either side of a screen with the animal tasked to move the stimulus to the center by moving the wheel (Fig. 3a). Serotonin stimulation was started at visual stimulus onset and lasted until the mice concluded their response or one second had passed, whichever came first. During a session, the prior probability of the stimulus appearing on the left or on the right varied in blocks of trials between 80% chance to appear on the left (80:20) or the right (20:80). Serotonin was stimulated in blocks of trials which were completely independent from the prior probability blocks. This way the mouse could not infer anything about the prior probability block structure from which trials were 5-HT stimulated.

**Figure 3.**
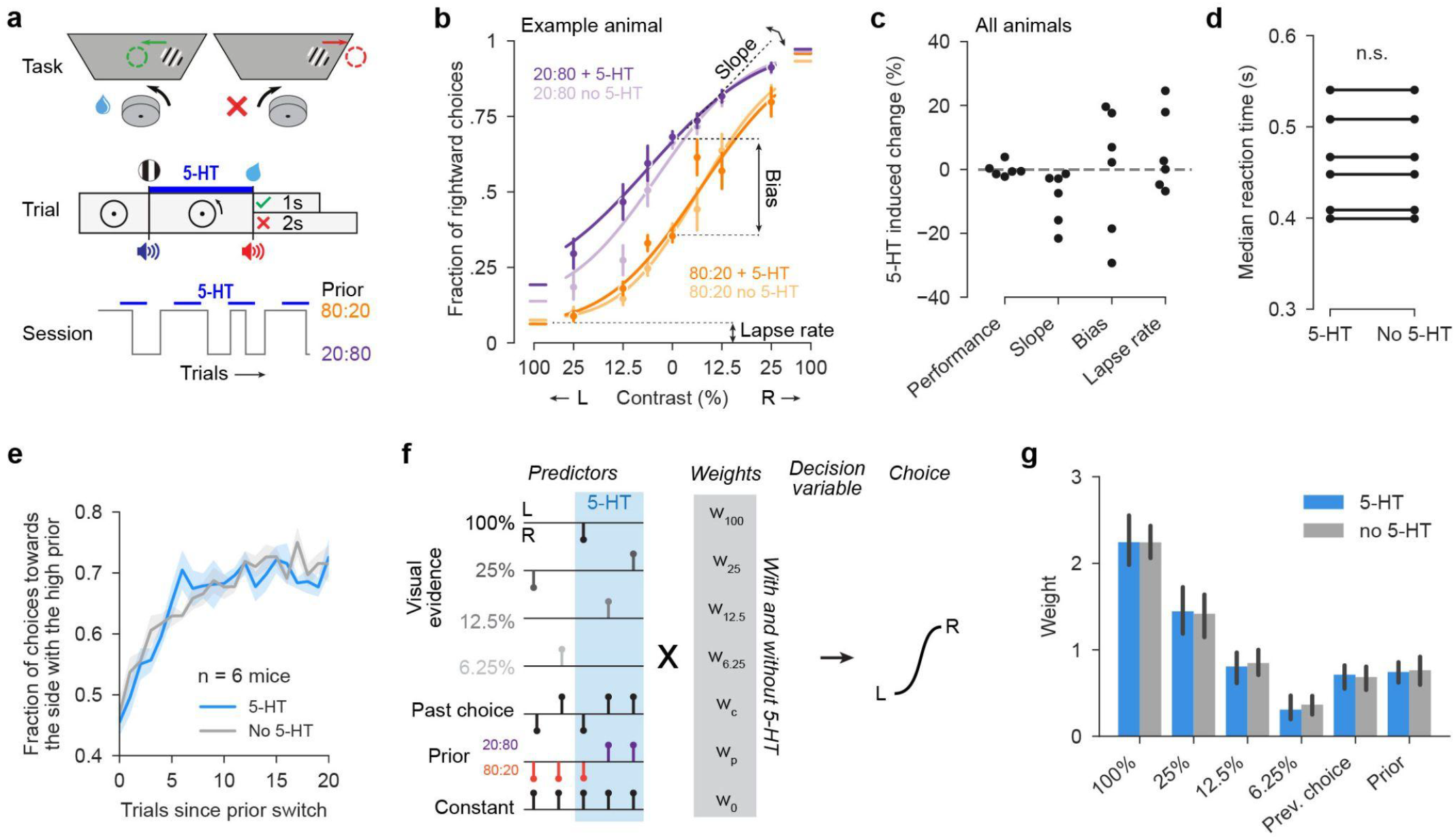
Serotonin stimulation does not affect decision making. **(a)** In the steering wheel task, mice are trained to position a stimulus in the center of the screen by turning a wheel. Optogenetic 5-HT stimulation is on from stimulus onset until the stimulus is either in the center (correct choice) or outside of the screen (incorrect choice). In blocks of trials there is a high probability of the stimulus appearing on the left (80% chance: 80:20) or the right (20:80). 5-HT is stimulated in blocks of trials which are independent from the prior blocks. **(b)** Psychometric curves of an example animal for 20:80 and 80:20 blocks, separated by whether 5-HT was stimulated. **(c)** The percentage change between 5-HT trials and non-stimulated trials for performance (percentage correct), slope, bias and lapse rate. Dots are animals. None are significantly different from zero (*p* > 0.05, t-test against 0). **(d)** The median reaction time for 5-HT stimulated and non-stimulated trials is not significantly different (paired t-test). Lines are animals. **(e)** Mice did not change their prior updating speed when stimulating 5-HT. The fraction of choices towards the side which had 80% chance of stimuli appearing did not increase more rapidly after a prior block switch. **(f)** Illustration of the design matrix of the probabilistic choice model fitted to the behavior, each column of stems is an example trial. Upward pointing stems indicate a +1 and downward pointing a -1 in the design matrix. Each predictor was inputted separately depending on whether the trial was 5-HT stimulated or not resulting in 13 individual predictors. **(g)** Weights of the predictors, bars are the mean and errorbar the s.e.m. over mice. For none of the pairs of predictors (5-HT and no 5-HT) there was a significant difference (*p* > 0.05, paired t-test).

Psychometric curves were constructed by plotting the percentage of rightward choices versus the visual contrast and fitting an error function. This was done separately for the prior probability blocks and the 5-HT stimulation blocks of trials. From the psychometric curves several metrics could be extracted: the slope (visual acuity), lapse rate (motivation), and bias (influence of the prior). We hypothesized that serotonin stimulation might impact the motivation of the mice and/or their dependence on the prior, therefore we expected to see differences in the lapse rate or the bias. However, we found no differences for any of the psychometric measures when comparing 5-HT stimulated and non-stimulated trials (Fig. 3c). There was also no difference in overall performance (% correct). Changes in motivation could also manifest themselves in a change in reaction times, however, we found no difference in reaction times (Fig. 3d).

Since serotonin is thought to be a surprise signal one would expect that in the 5-HT stimulated blocks of trials mice are more flexible; if everything is surprising the optimal behavior is to rapidly update your priors. To see if this was the case in our task, we focussed on trials where the prior probability switched from one side to the other. If mice are more flexible we expected to see that they would more rapidly update the bias in their choices towards the side that the prior probability switched, e.g. if the prior switched from 80% chance on the left to 80% chance on the right mice would start preferring the right side more quickly after a prior switch. However, we found no difference in the speed at which mice updated their response bias after a prior switch (Fig. 3e).

It could still be the case that 5-HT stimulation had a more subtle effect on the choices of the animals; maybe 5-HT only affected the choices at certain visual contrast levels for example. To investigate this possibility we fitted a probabilistic choice model^26,27^ to the behavior with predictors for visual evidence, the past choice, and the stimulus prior, which were split by 5-HT stimulation (Fig. 3f). However, there was no difference in the weight for the 5-HT stimulated versus the unstimulated trials for any of the predictors (Fig. 3g). Taken together these results show that, in this task, 5-HT stimulation does not affect usage of the prior, motivation, flexibility, or choices in general.

### Serotonergic modulation of neural dynamics during task performance

Although we found no effect of 5-HT stimulation on the behavior of the animals, at the neuronal level we observed substantial serotonergic modulation. Similarly to the modulatory effects during quiet wakefulness, serotonin stimulation modulated the activity of single neurons both positively and negatively (Fig. 4a). The percentage of modulated neurons across the brain was, depending on the brain region, between 20 and 40% (Fig. 4b). One notable exception was the amygdala which contained around 5% significantly modulated neurons. The modulation index across regions showed a similar pattern compared to 5-HT modulation during quiet wakefulness (Fig. 4c). Overall the strength of the serotonergic modulation was weaker during task performance compared to quiet wakefulness, brain regions showed significantly less 5-HT modulated neurons during the task compared to quiet wakefulness (Fig. 4d). In short, during task performance there is widespread 5-HT modulation of neural dynamics across the brain, albeit less strong than during quiet wakefulness.

**Figure 4.**
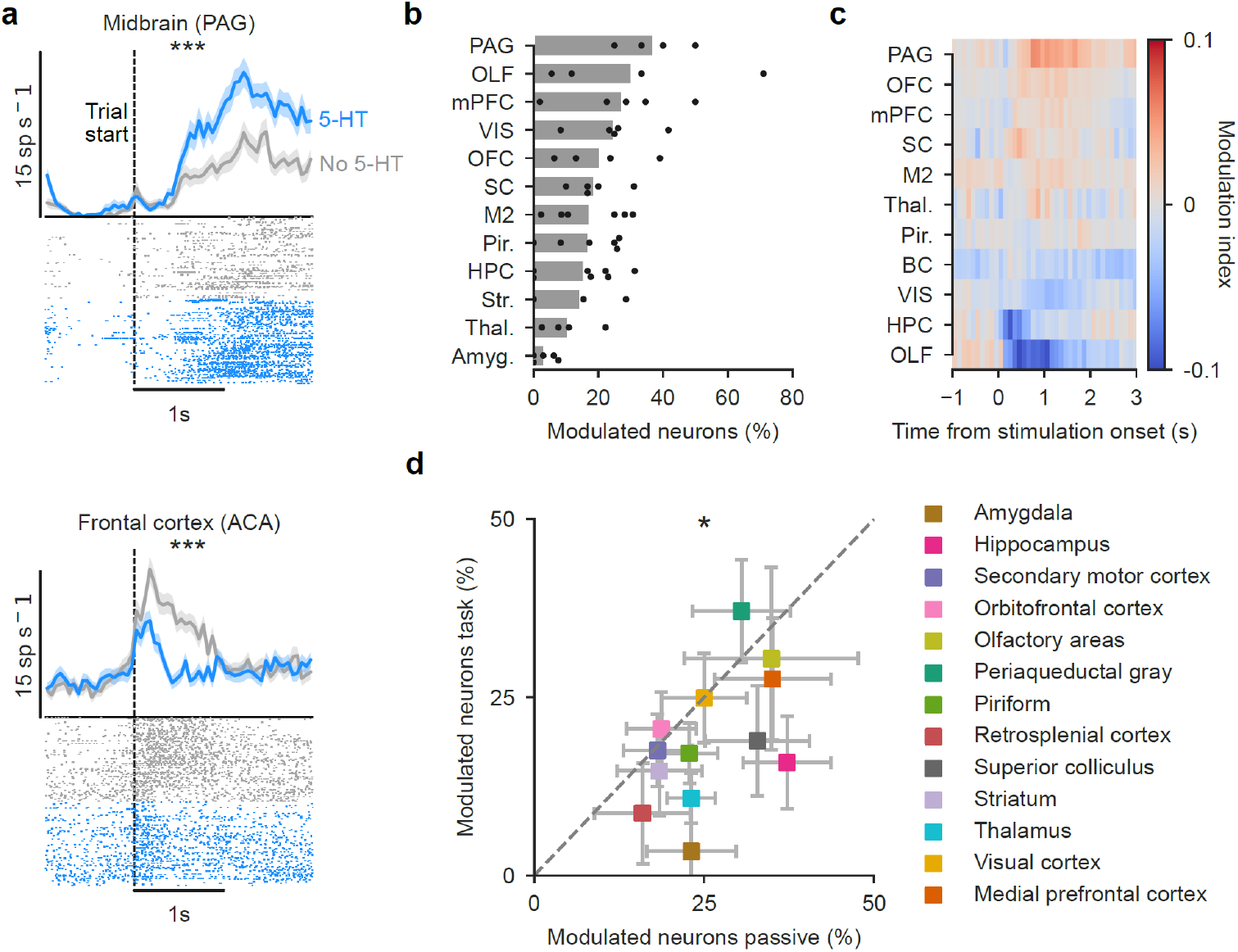
Serotonergic modulation of neural dynamics during task performance. **(a)** Two typical example neurons which show significant 5-HT modulation during task performance. Neural activity is conditioned to the start of the trail: visual stimulus onset with (blue line) or without (grey line) 5-HT stimulation. Thick lines indicate the mean firing rate over trials (50 ms bins with 25 ms Gaussian smoothing window), with shading indicating the standard error of the mean. The bottom part of the plot shows the spike raster in which each row is a trial with ticks showing individual spikes. ***: two-sample ZETA test *p* < 0.001. **(b)** The percentage of significantly modulated neurons (*p* < 0.05, two-sample ZETA test between 5-HT stimulated and unstimulated trials) per region. Black dots are individual mice and grey bars the mean over mice (sert-cre only). **(c)** The modulation index (red is positive and blue is negative 5-HT modulation) per region. **(d)** Significantly fewer neurons were 5-HT modulated during task performance versus passive stimulation. Colored squares are the mean per region, grey whiskers indicate s.e.m. * *p* = 0.012, paired t-test.

### Serotonin modulates neural dynamics in subspace orthogonal to the choice vector

How is it possible to observe widespread 5-HT modulation in the brain but no effect on behavior? One possible explanation is that the geometry of the serotonergic modulation was such that it did not interfere with the neural code for the choice of the animal. To investigate this, we performed a manifold analysis (as in ref. ^28^) by taking the trial-averaged peri-event time histogram of all neurons, across all sessions and mice, and combining them into a single super-session. Neural data was centered at the moment the mouse initiated the response, defined as the time of the first movement of the wheel after stimulus onset. When splitting trial by choice, irrespective of 5-HT stimulation, the trajectories for left and right choices diverged over time leading up to the choice moment (Fig. 5a). This was quantified by computing the Eucledian distance between the trajectories at each timepoint in neural state space. The choice separation indeed increased over time, leading up to the choice, and was significantly larger than the null-distribution (Fig. 5b). When performing the manifold analysis separately per brain region and analyzing the last time point before the choice moment, we found that the following brain regions showed a significant choice separation: supplementary motor cortex, medial prefrontal cortex, orbitofrontal cortex, superior colliculus, hippocampus and visual cortex (Fig. 5c).

**Figure 5.**
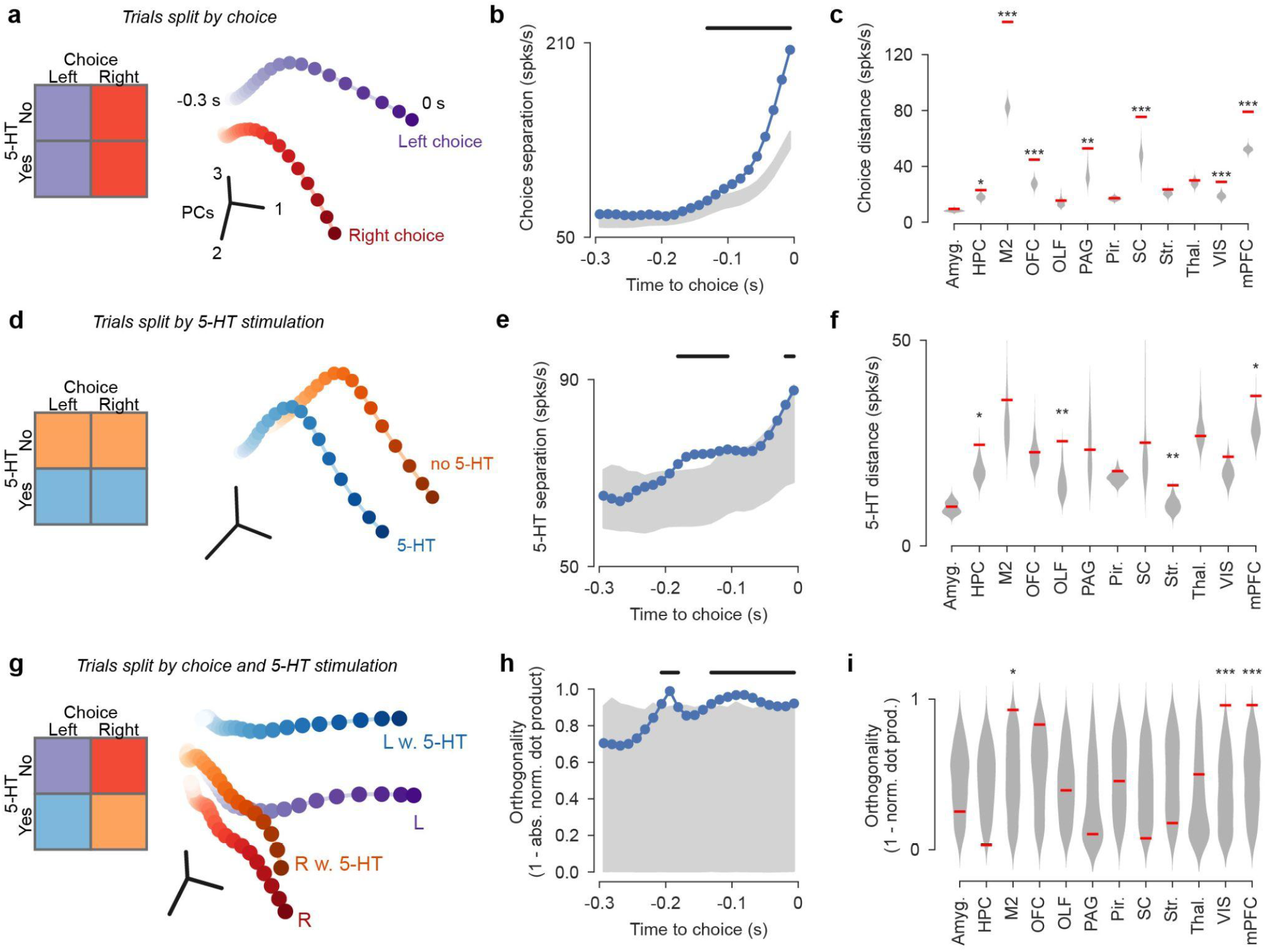
Serotonin modulates neural dynamics in a subspace orthogonal to the choice vector. **(a)** PCA projection of trial-averaged activity per neuron centered at the choice (t0 = first movement of the wheel), trials were split into left and right choices, regardless of 5-HT stimulation. **(b)** The separation between left and right choices, defined as the Eucledian distance in neural space between the population vectors at each timepoint. Grey area represents the 95% confidence interval of the null-distribution obtained by shuffling the choices. Black bar above the plot indicates significant timepoints (*p* < 0.05). **(c)** Choice separation for the last time point before the choice, computed per brain region. Red dashes indicate the choice separation on the original dataset, grey violins are the shuffled null-distribution. * *p* < 0.05, ** *p* < 0.01, *** *p* < 0.001; permutation testing. **(d)** PCA projections as in (a) but for splitting the trials into 5-HT stimulated and non-stimulated trials, regardless of choice. **(e)** Separation between 5-HT stimulated and non-stimulated trials. Grey area indicates 95% confidence interval of null-distribution, obtained by generating pseudo 5-HT stimulation blocks. **(f)** 5-HT separation for the last time point before the choice per brain region as in (c). **(g)** PCA trajectories of left and right choices split by 5-HT stimulation. **(h)** The orthogonality between the average choice axis and the stimulation effect axis in (g), computed per time point. **(i)** The orthogonality of the last time point before the choice calculated per brain region.

We next computed the 5-HT separation by splitting the trials by serotonin stimulated and unstimulated trials, regardless of choice. We found two distinct trajectories in PCA space for the trials with and without 5-HT stimulation (Fig. 5d). These trajectories similarly diverged over time, leading up to the choice, but less so compared to the choice separation (Fig. 5e). The magnitude of the separation was also less strong for the 5-HT versus the choice, note the difference in scaling of the y-axes. Looking at individual regions, we found that 5-HT separation, at the last time point, was significant in striatum, olfactory areas, hippocampus and medial prefrontal cortex (Fig. 5f).

Finally, we constructed trajectories for left and right choices, split by 5-HT stimulation (Fig. 5g). We observed from the PCA trajectories that the trajectories with 5-HT stimulation (blue and orange in Fig. 5g) were situated on the orthogonal axis compared to the axis between the choices without 5-HT stimulation (red and purple in Fig. 5g). To quantify this, we calculated how orthogonal the stimulus effect vector was from the average choice vector. We found that the orthogonality between these vectors increased as the choice moment approached and was significantly more orthogonal than the shuffled null-distribution (Fig. 5h). When doing this analysis for the last time point before the choice per region, we found significant orthogonal geometry between the choice and the stimulation in medial prefrontal cortex, visual cortex and supplementary motor area (Fig. 5i).

To conclude, both choice and 5-HT stimulation could be separated by manifold trajectories leading up to the choice moment. We found that the geometry between choice and 5-HT trajectories was such that the stimulation axis was orthogonal compared to the choice axis. This suggests a mechanism by which serotonin can exert a modulatory influence on task representations without hampering goal-directed behavior itself.

## Discussion

In this study we found that serotonin stimulation induces strong effects on spontaneous behaviors during quiet wakefulness. Also at the neuronal level, serotonin modulated neural dynamics across all recorded brain regions. It was surprising, therefore, that we did not find any effect of 5-HT stimulation on mouse behavior during active engagement in a decision-making task. Though, we still observed widespread modulation of neural dynamics, albeit weaker compared to during quiet wakefulness. How can serotonin induce widespread modulation of neural activity while not affecting goal-directed behavior? Manifold analysis revealed that serotonergic modulation of neural dynamics was confined to a subspace which was orthogonal compared to the choice axis.

At the neuronal level, reports on how serotonin stimulation influences activity have been conflicting. Electrophysiological recordings in the visual cortex during visual stimulation^17^ and in the piriform cortex during olfactory stimulation^19^ found that serotonin stimulation suppressed neural activity. Also using fMRI, serotonin was found to have a brain-wide inhibitory effect^18^. A more recent fMRI study, however, found brain-wide excitatory effects of 5-HT stimulation^8^ and posits that previous studies had been done under anesthesia (although note that ref. ^17^ included awake experiments), which explains the differences in findings. Here we did large-scale electrophysiological recordings across the brain during 5-HT stimulation in the awake condition and found a more nuanced picture. While the periaqueductal gray is excited and the hippocampus is suppressed, most regions showed a balance between excitation and inhibition. Even the regions that were suppressed included individual neurons that were excited and vice versa. We therefore conclude that serotonin does not induce brainwide excitation or inhibition but instead bidirectionally modulates neural dynamics across the brain.

Previous studies found that optogenetic stimulation of serotonin release resulted in increased persistence^13,14,29–31^. This led us to hypothesize that, in our task, we would find that 5-HT stimulation altered the way in which mice incorporated prior knowledge with current sensory evidence, or the speed by which they updated their prior. Contrary to our hypothesis, we found no evidence for either of these two things. One possibility is that the serotonin stimulation was not started early enough to influence the decision, serotonin is known to be quite slow. However, we found that the onset of 5-HT modulation could be as early as 25 ms after stimulation onset (in hippocampus) while reaction times were typically around 400-500 ms. Furthermore, serotonin was stimulated in blocks of trials so even if the stimulation was too late to impact the decision on that trial, it could still influence how the prior was updated, a process which happens over several trials.

We stimulated serotonin release by activating serotonergic neurons in the dorsal raphe nucleus and measured its effect by recording neural dynamics in the rest of the brain. The observed effects of 5-HT stimulation on neural dynamics, however, cannot be solely attributed to direct serotonergic input to the recorded neural circuits. Serotonin broadcasts all across the brain resulting in direct projections, but also myriad indirect pathways, to the neural population under scrutiny. Therefore, the observed modulation of neural dynamics is most likely a mixture of direct serotonergic modulation and indirect modulation through different pathways. For example, serotonin might modulate subcortical areas in the brain related to arousal, which in turn modulate neural activity in cortex. A notable example is the hippocampus, we observed strong and rapid inhibition of the hippocampus. However, there is no strong direct projection from the dorsal raphe nucleus to the hippocampus. A likely explanation is that serotonergic dorsal raphe neurons project to the median raphe which in turn projects to the hippocampus.

We observed that there were fewer significantly 5-HT modulated neurons in the task compared to the passive condition. This could indicate that the serotonergic input is weaker during active behavior but there are two alternative explanations. Firstly, it’s simply easier to establish significance in the passive condition since the baseline firing rates are relatively low. Secondly, in the passive condition, neurons are spontaneously active around the optimum of their dynamic range^32^, any modulatory input at this time will have a relatively strong effect on the firing rate. During behavior, neurons receive task-induced input which drives their firing rate away from their optimal level, reducing the relative effect of a neuromodulatory input. It is therefore possible that optogenetically induced serotonergic input is of similar strength during quiet wakefulness and task performance but because of the aforementioned reasons its effect on neural activity is weaker during behavior.

Our main finding is that the modulatory effect of serotonin is confined to a subspace which is orthogonal to the choice axis. The medial prefrontal cortex, for example, strongly encodes the upcoming choice but is also modulated by serotonergic input. By orthogonalizing the choice and the 5-HT axis, downstream regions of the mPFC can either read out decision-related information, by taking a linear weighted sum of their inputs and projecting the activity onto the choice subspace, or they can project their inputs along the 5-HT axis to extract neuromodulatory information. This hints to the existence of state-dependent “neuromodulatory subspaces”; extra dimensions that allow neuromodulatory systems to modulate information potentially to create new, orthogonalized, representations of the environment to which different rules can be applied. This neuromodulatory subspace might serve functions such as the relaying of information regarding internal state. Another possibility is that the orthogonal subspace allows for the rapid encoding of new features of the environment which aids flexible behavior. When the environment changes this results in a surprise response which drives serotonin release^3^. Subsequently, serotonin modulates the neural dynamics into the neuromodulatory subspace, allowing the brain to distinguish the novel situation from the old one. Hypothetically, this expansion of the neural code facilitates the rapid updating of neural representations to match changes in the environment, and ultimately aid flexible behavior.

## Materials and Methods

### Animal subjects

All procedures were approved by the Portuguese Veterinary General Board. Seventeen C57Bl/6 mice (7 males and 10 females) were used in this study. Of these, 11 were SERT-cre and 6 were wild-type control mice.

### Headbar & fiber implantation and virus injection surgery

The surgery was performed in a robotic stereotactic apparatus (Neurostar) equipped with an automatic Hamilton syringe. Mice were anesthetized using 3% isoflurane (by volume in O2) which was lowered during the surgery to 1.5-2%. Mice were fixed to the stereotactic frame using cheek bars and placed on a heating pad. Their eyes were covered with eye ointment (Vitaminoftalmina A, DÁVI - Farmacêutica) and protected against the surgical lights with aluminum foil. Analgesia was administered after anesthesia induction by a subcutaneous injection of 5 mg/kg Carprofen. The head was shaved using clippers and any remaining hair was removed using depilatory cream (Veet). The skin was disinfected using betadine and ethanol and an incision was made along the dorsal skull. The periost was carefully removed from the skull and the skull was cleaned with a bone scraper. The tendons that attach to the skull on the posterior sides were severed to create enough space for the headbar. The muscles attaching to the posterior skull plate were detached from the skull and the skin was glued to the skull using Vetbond (3M). The skull was aligned by touching the skull with the glass capillary of the Hamilton syringe and adjusting the cheek bar and nose clamp height until the skull was level (0.04 mm tolerance). Two injections of AAV2/1.EF1a.double floxed ChR2.EYFP.WRPE.HGHpA were made at an angle of 32° at -6.54 and -6.74 mm AP and 0 mm ML relative to Bregma ^33^. The injections were made at a depth of 4.02 mm, the motorized injection system was set up so that the pipette would be pulled upwards for 0.3 mm over the course of the injection. The two injections were both 150-250 nl in volume and the injection speed was such that each injection took 10 minutes. An optical fiber (MFC 200/230-0.048 4mm ZF1.25(G) FLT, Doric) was implanted at a 32° angle, at -6.64 mm AP, 0 mm ML, 3.77 mm DV. To keep a large area of the skull accessible for Neuropixel recordings, the headbar was implanted over the optical fiber by placing the fiber through a hole in the headbar itself. Because of the large angle at which the fiber was implanted, the headbar had to be angled 5-10° to allow the fiber to fit through the hole in the headbar. This was achieved by placing the headbar in a 3D printed holder which was attached to the stereotactic arm. The headbar was cemented to the skull using C&B Superbond (Sun Medical), of which a layer was also applied over the surface of the skull to prevent infections. All exposed areas were covered up with black dental cement (Contemporary Ortho-Jet, MediMark) which was also used to create a well around the skull surface. Lastly, a layer of UV glue (Optical Adhesive 61, Norland) was used to fill up the well which was cured using a UV flashlight (UV301D, LightFe). Animals were allowed to recover for a week after this surgery.

### Craniotomy surgery

Anesthesia, analgesia, eye covering, and heating of the body was the same as the implantation surgery. The headbar was fixed in a 3D printed headbar holder which replaced the position of the cheek bars in the stereotact. The layer of optical glue was removed using tweezers and the layer of C&B Superbond was drilled away using a dental drill. Three craniotomies were made in the left hemisphere at: (1) -1.1 mm ML, 2 mm AP; (2) -2.2 mm ML, -2 mm AP; (3) -1 mm ML, -3.3 mm AP relative to Bregma. The craniotomies were circular with an inner diameter of ∼0.5 mm. They were made using a microdrill (78001, RWD Life Science) which was equipped with a 0.3 mm drill bit (122-2008, Gesswein). The exposed brain was covered with duragel (DOWSIL 3-4680, Dow Corning) which protects the brain and allows probe insertions over multiple days^34^. A ground pin was implanted in the right hemisphere at around 2 mm ML, -1 mm AP, the pin was cut short such that it would rest on top of the brain surface without causing substantial damage to the cortical surface. Lastly, a protective cap was placed on top of the well and fixed in place with Kwik-Cast (World Precision Instruments).

### Neuropixel recording rig

The rig consisted of a 750 × 900 mm airtable (M-VIS3036-SG2-325A, Newport) on which a sound attenuated chamber was built. In the center of the airtable there was a holder for the mouse body with clamps on either side to fixate the headbar. Furthermore, the rig was outfitted with components to allow head-fixed behavior: a steering wheel, iPad screen, and speaker. These components were not used for the purpose of this manuscript, the steering wheel could move but did not do anything and the iPad screen was on but only showed a gray screen. Three cameras (CM3-U3-13Y3M-CS, FLIR PointGrey with MVL16M23, Thorlabs lenses) were positioned around the body holder, two pointing to either side of the mouse’s head and one looking down on the body from the top. Two motorized micromanipulators (uMp-4, Sensapex) were positioned on either side of the mouse holder to allow simultaneous insertion of two Neuropixel probes. Electrophysiological signals were recorded using a PXI system (PXIe-1071, PXIe-8381, PXIe-6341, National Instruments) with a BNC breakout board (BNC-2210, National Instruments) to synchronize neural data with optogenetics and video. Light stimulation was delivered by an LED (CLED 465 mm, Doric) which was powered by a programmable open-source LED-driver^35^ (Cyclops, Open Ephys). The LED-driver was started and stopped by a sequence of TTL pulses emitted by a Bpod (Sanworks).

### Behavioral task

The behavioral task was the International Brain Lab steering wheel task which has been described in detail in ref. ^27^, the procedures for handling and training mice in the task are described in ref. ^36^. Briefly, head-fixed mice were tasked to position a Gabor stimulus, which appears either on the left or right side of the screen, in the center of the screen by moving a steering wheel. A trial started with a quiescent period of 400-700 ms in which the mouse should keep the wheel still, movement of the wheel resulted in the rest of the quiescent period. After this, the visual stimulus with a contrast of [100%, 25%, 12.5%, 6.25% or 0%] appeared on the screen accompanied with an auditory cue. Movement of the wheel to position the stimulus in the center of the screen was rewarded with 3 μl of 10% sucrose water followed by a 1s inter-trial interval, moving the stimulus outside of the screen was punished with a white noise burst and a 2s inter-trial interval. The prior probability that the stimulus would appear on the left or the right was varied in blocks of trials, each block was randomly varied in length between 20 and 100 trials by drawing from a truncated geometric distribution (average block length was 51 trials). In this manuscript, we used the task exactly as described in ref. ^37^ with one change: there was no block of 90 trials at the beginning of each session in which the prior probability was 50/50.

### Optogenetic stimulation

Light intensity was 5 mW as measured at the tip of the optical fiber during continuous illumination. The stimulation pulse train consisted of 10 ms pulses at 25 Hz. To reduce light-induced electrical artifacts, the Cyclops LED-driver was programmed to deliver 10 ms light pulses which contained a 1 ms cosine ramp on either side such that the light intensity was varied smoothly^38^. During the behavioral task, light stimulation was on from the moment of stimulus onset until the moment the mouse concluded their choice, or after one second had elapsed, whichever came first. 5-HT was stimulated in blocks of trials, the length of a stimulated block was randomly drawn from a truncated geometric distribution (between 20 and 100 trials, average 51). Contrary to the prior blocks a fixed block length was used per session, randomization of the block length occurred only between sessions. Passive stimulation during quiet wakefulness consisted of 50 repetitions of 1s pulse trains with an inter-stimulation interval drawn from a truncated exponential distribution with a factor of 7s, capped at a minimum of 3s and a maximum of 13s.

### Neuropixel recordings

To allow post-hoc reconstruction of the probe tract, the tip of the probe was lowered into a small droplet of CellTracker CM Di-I (Termo-Fisher Scientific) using the micro-manipulator^39^. The mouse was head-fixated in the apparatus and the protective cap was removed. A split-wire was inserted into the ground pin and attached to the grounds of the two Neuropixel headstages. Both probes were zeroed on Bregma and inserted in their respective recording targets for that session. For full details regarding the procedure see refs. ^37,40^. The targets were four fixed pairs of insertion coordinates relative to Bregma, recording sessions were randomized over mice.

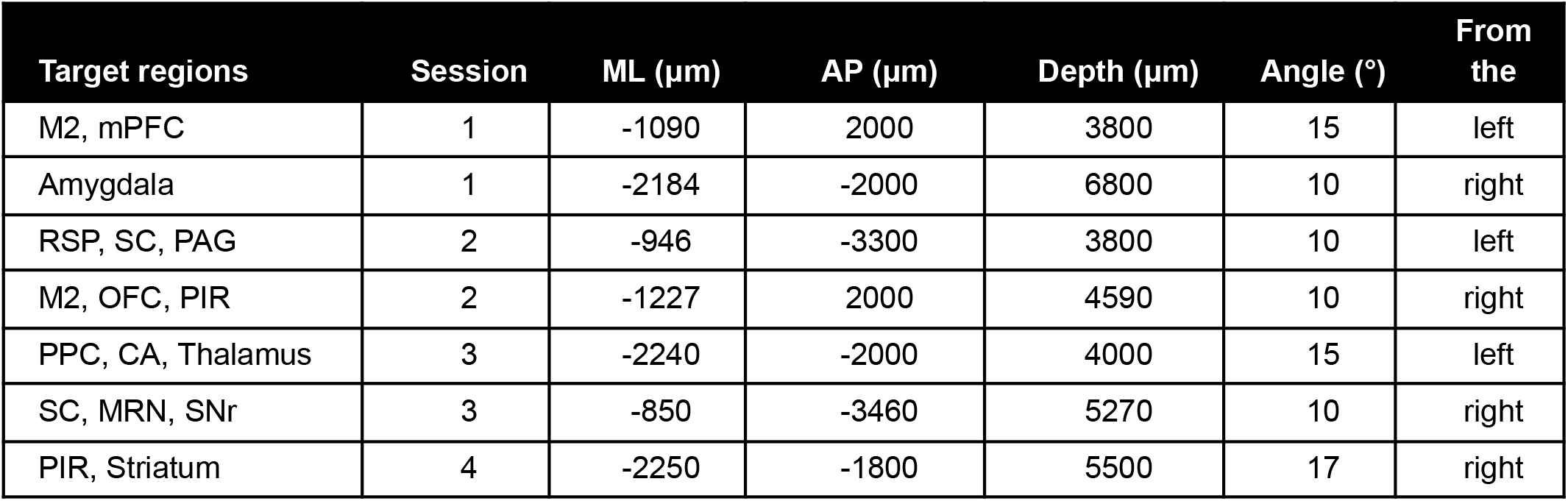

| Target regions | Session | ML (μm) | AP (μm) | Depth (μm) | Angle (°) | From the |
| --- | --- | --- | --- | --- | --- | --- |
| M2, mPFC | 1 | -1090 | 2000 | 3800 | 15 | left |
| Amygdala | 1 | -2184 | -2000 | 6800 | 10 | right |
| RSP, SC, PAG | 2 | -946 | -3300 | 3800 | 10 | left |
| M2, OFC, PIR | 2 | -1227 | 2000 | 4590 | 10 | right |
| PPC, CA, Thalamus | 3 | -2240 | -2000 | 4000 | 15 | left |
| SC, MRN, SNr | 3 | -850 | -3460 | 5270 | 10 | right |
| PIR, Striatum | 4 | -2250 | -1800 | 5500 | 17 | right |

### Pre-processing and spike sorting

The fully automated pipeline of the IBL was used for pre-processing and spike sorting of the raw Neuropixel data^24^. As a spike sorting algorithm, the open-source Python implementation of Kilosort was used, which was developed by the IBL (pykilosort; https://github.com/int-brain-lab/pykilosort).

### Histology

After all recordings were done the mouse was perfused and the fixed brain tissue was shipped to the Sainsbury Wellcome Centre (Londen, United Kingdom), according to IBL protocol^41^. There, whole-brain reconstruction was done using serial-sectioning two-photon imaging^42^. After registration of the full brain volume to the Allen Brain Atlas, the fluorescent probe tracts were manually traced in the 3D volume using LASAGNA^43^. Finally, the electrophysiological signatures along the probe were manually aligned to the brain regions along the histological tract with the IBL ephys alignment GUI.

### Data analysis

All analysis was performed in Python using the ONE protocol to access the data, all the code to reproduce the figures is available: https://github.com/guidomeijer/SerotoninStimulation/.

### Pupil size, sniffing and whisking

Behavioral markers were extracted from the face video. Pupil size was defined as a circle fitted to the four DeepLabCut (DLC)^44^ key points surrounding the pupil, for full details regarding the camera setup and DLC pipeline see ref. ^45^. Sniffing was estimated by taking the frame-to-frame displacement of the tracked key point on the tip of the nose. Whisking was estimated by calculating the movement energy in an ROI around the whisker pad. Pupil size, sniffing, and whisking traces were subsequently smoothed with a third-order Savitzy-Golay filter with a window size of 61 for pupil size and 31 for sniffing and whisking. Missing data in the pupil size trace due to eye closure or obstruction of the view of the pupil were interpolated. To account for inter-animal and session-to-session variation, relative changes were computed by calculating the percentage change to baseline for each session independently; baseline was defined as the 2nd percentile of all values of the session^20^.

### Hippocampal sharp wave ripple detection

From all Neuropixel insertions in the dorsal hippocampus the channels in CA1 were selected. Sharp wave ripples were detected on the recording channel which was in the CA1 pyramidal cell layer, as defined by the channel with the highest AP band RMS. The LFP trace from this channel was band-pass filtered for the ripple band (150-250 Hz) using a fifth-order Butterworth filter. Sharp wave ripples were detected as deflections above six standard deviations with a minimum time interval between them of 100 ms.

### Neural responsiveness, modulation, and latency

To determine whether a neuron was significantly modulated by 5-HT stimulation, the ZETA-test for neural responsiveness was used^46,47^, specifically the Python implementation *zetapy*. This test makes no assumptions and can detect complex multi-phasic response patterns, such as the ones often observed in response to serotonergic input. Furthermore, the ZETA test defines the temporal onset of the modulation as the first crossing of the half-height of the deviation from baseline.

The direction and magnitude of serotonergic modulation was quantified with a modulation index^19^, which was calculated using receiver operator characteristics (ROC) analysis. Per neuron, the spike counts in a reference time window were compared to a target time window. The area under the ROC curve (auROC) was calculated which resulted in a value in the range of [0, 1]. To center this quantity at zero it was recalculated as follows:

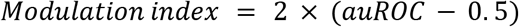

Which resulted in an index in the range of [-1, 1] whereby negative values reflect decreases in firing rates, during stimulation compared to baseline, and positive values reflect increases.

### Psychometric curve fitting

Visual contrast values were signed such that contrasts presented on the left of the screen were negative and contrasts on the right positive. The percentage of times the mouse chose the right stimulus was plotted on the y-axis and an error function with two lapse rates, one for each side (γ & λ), a bias term (μ), and a slope parameter (σ) was fitted through these points using the python package psychofit following the methods in ref. ^37^.

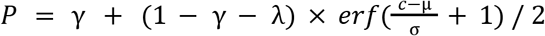

Here *P* is the probability of choosing the right side and *c* is the signed visual stimulus contrast. The slope parameter (σ) is defined here as a scaling of the horizontal axis such that larger values translate into shallower slopes. This is counter-intuitive with respect to how slopes are typically defined (larger values mean steeper slopes), therefore the slope was transformed as follows: δ = 1/δ.

### Probabilistic choice model

To investigate the impact of serotonin stimulation on the choices of the animal, a probabilistic choice model was fitted to the behavior^26,37^. The model is a binomial logistic regression model where the probability of choosing left or right is estimated from sensory and non-sensory predictors. A separate set of predictors was inputted into the model for 5-HT stimulated and non-stimulated trials. Probabilities were obtained from the logistic transformation of decision variable *z*, which was the result of a weighted linear sum of all the predictors *W*. For each trial *i, z* is calculated as follows:

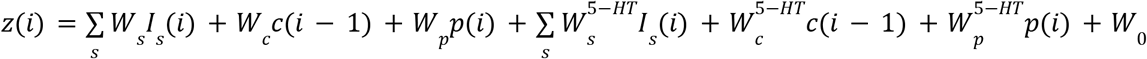

*W*_*s*_ is the weight for each stimulus contrast *s* ∈ {6.25, 12.5, 25, 100} with *I*_*s*_ indicating a -1 if the stimulus appeared on the left and a +1 if it appeared on the right in trial *i*.

*W*_*c*_ is the weight for the previous choice: -1 if the mouse chose left on the previous trial and +1 if it chose right.

*W*_*p*_ is the weight for the prior probability of the stimulus appearing left or right; -1 if there is an 80% chance of the stimulus appearing left and +1 if there is an 80% chance of it appearing right.

*W*_*0*_ is a constant term.

If trial *i* is a 5-HT-stimulated trial, all the corresponding terms are passed to the *W*^*5-HT*^ weights while passing a 0 to the other weights, and vice versa. A design matrix was constructed this way for all the trials of an animal concatenated together over sessions. Subsequently, the model was fitted with regularized maximum likelihood estimation as implemented by the *Logit*.*fit_regularized* function from statsmodels^48^.

#### Population manifold analysis

The same procedure was applied as in ref. ^28^, briefly: for each neuron a peri-event time histogram (PETH) was constructed by centering the spike times to either the start of the wheel movement (Fig. 5) or the onset of the visual stimulus (Supp. Fig. 5). Spike trains were binned in 12.5 ms bins with a 20 ms Gaussian smoothing kernel and the mean over trials was computed. Trials with a reaction time of < 0.1s were excluded because the 5-HT stimulation started with visual cue onset and could probably not have affected the choice in this time. Also trials with reaction times of >1.2s were excluded since the 5-HT stimulation only lasted for a maximum of 1s. The resulting trials were then split in several different ways: by choice (left and right), 5-HT stimulation (stimulated trials and unstimulated trials), or both (left choice with 5-HT, left choice without 5-HT, right choice with 5-HT, and right choice without 5-HT). The resulting PETHs of all neurons, across all sessions and mice, were stacked into one super-session. From this super-session choice separation was computed, at each timepoint, as the Eucledian distance in neural space between the trajectories for the left versus the right choices. 5-HT separation was the Eucledian distance between the 5-HT stimulated and unstimulated trajectories.

Before applying PCA, a normalization step was performed by mean-subtracting each PETH. Subsequently, PCA was performed to reduce the n-dimensional matrix from *n* neurons to three principal components. Orthogonality between the choice axis and the stimulation axis was determined by first computing the average direction of the choice axis:

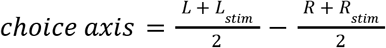

Here L is the coordinate of the left choices without 5-HT stimulation in 3-dimensional PCA space, R the right choice and L_stim_ and R_stim_ are the left and right choices with 5-HT. Essentially this means that the choice axis is defined as the vector between the average of the left and right choices, regardless of stimulation. The 5-HT effect axis was determined as follows:

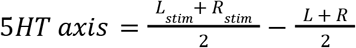

Subsequently, the vectors were normalized and the dot product was calculated between them. The original dot product is such that smaller values indicate a more orthogonal angle, to make this value easier to interpret the final orthogonality measure as defined as *1 -* |*dot product*|. This resulted in a measure which is scaled between 0 and 1 in which higher values indicate more orthogonal angles.

Generating null-distributions is not straightforward because of the block design of the task and the 5-HT stimulation, if not done properly one might inflate the effect because of within-block temporal autocorrelations^49^. For the choice separation it is insufficient to simply shuffle the choices over trials since the choice of the animal is strongly influenced by the prior probability of the stimulus side, which varies in blocks of trials. To account for this issue, choices were shuffled separately within each stimulus prior block. The 5-HT stimulation was administered in blocks of trials, posing the same problem. Therefore, instead of shuffling the stimulated trials, pseudo-blocks were generated in the same way the original blocks were.

## Data and code availability

All the data in this manuscript is publicly available through the Open Neurophysiology Environment (ONE) protocol: https://int-brain-lab.github.io/ONE/. All the code used to generate the figures in this manuscript can be found on Github: https://github.com/guidomeijer/SerotoninStimulation.

## References

1. Clarke, H. F., Dalley, J. W., Crofts, H. S., Robbins, T. W. & Roberts, A. C. Cognitive Inflexibility After Prefrontal Serotonin Depletion. Science 304, 878–880 (2004).

2. Grossman, C. D., Bari, B. A. & Cohen, J. Y. Serotonin neurons modulate learning rate through uncertainty. Curr. Biol. 32, 586-599.e7 (2022).

3. Matias, S., Lottem, E., Dugué, G. P. & Mainen, Z. F. Activity patterns of serotonin neurons underlying cognitive flexibility. eLife 6, e20552 (2017).

4. Ligneul, R. & Mainen, Z. F. Serotonin. Curr. Biol. 33, R1216–R1221 (2023).

5. Azmitia, E. C. & Segal, M. An autoradiographic analysis of the differential ascending projections of the dorsal and median raphe nuclei in the rat. J. Comp. Neurol. 179, 641–667 (1978).

6. Ren, J. et al. Single-cell transcriptomes and whole-brain projections of serotonin neurons in the mouse dorsal and median raphe nuclei. eLife 8, e49424 (2019).

7. Li, Y. et al. Serotonin neurons in the dorsal raphe nucleus encode reward signals. Nat. Commun. 7, 1–15 (2016).

8. Hamada, H. T. et al. Optogenetic activation of dorsal raphe serotonin neurons induces brain-wide activation. Nat. Commun. 15, 4152 (2024).

9. Zhou, J., Jia, C., Feng, Q., Bao, J. & Luo, M. Prospective Coding of Dorsal Raphe Reward Signals by the Orbitofrontal Cortex. J. Neurosci. 35, 2717–2730 (2015).

10. Cohen, J. Y., Amoroso, M. W. & Uchida, N. Serotonergic neurons signal reward and punishment on multiple timescales. eLife 4, e06346 (2015).

11. Harkin, E. F., Grossman, C. D., Cohen, J. Y., Béïque, J.-C. & Naud, R. A prospective code for value in the serotonin system. Nature 1–8 (2025).

12. Hyun, J. H., Hannan, P., Iwamoto, H., Blakely, R. D. & Kwon, H.-B. Serotonin in the orbitofrontal cortex enhances cognitive flexibility. 2023.03.09.531880 Preprint at 10.1101/2023.03.09.531880 (2023).

13. Miyazaki, K. W. et al. Optogenetic Activation of Dorsal Raphe Serotonin Neurons Enhances Patience for Future Rewards. Curr. Biol. 24, 2033–2040 (2014).

14. Lottem, E. et al. Activation of serotonin neurons promotes active persistence in a probabilistic foraging task. Nat. Commun. 9, 1000 (2018).

15. Correia, P. A. et al. Transient inhibition and long-term facilitation of locomotion by phasic optogenetic activation of serotonin neurons. eLife 6, e20975 (2017).

16. Ren, J. et al. Anatomically Defined and Functionally Distinct Dorsal Raphe Serotonin Sub-systems. Cell 175, 472-487.e20 (2018).

17. Azimi, Z. et al. Separable gain control of ongoing and evoked activity in the visual cortex by serotonergic input. eLife 9, e53552 (2020).

18. Grandjean, J. et al. A brain-wide functional map of the serotonergic responses to acute stress and fluoxetine. Nat. Commun. 10, 1–10 (2019).

19. Lottem, E., Lörincz, M. L. & Mainen, Z. F. Optogenetic Activation of Dorsal Raphe Serotonin Neurons Rapidly Inhibits Spontaneous But Not Odor-Evoked Activity in Olfactory Cortex. J. Neurosci. 36, 7–18 (2016).

20. Cazettes, F., Reato, D., Morais, J. P., Renart, A. & Mainen, Z. F. Phasic Activation of Dorsal Raphe Serotonergic Neurons Increases Pupil Size. Curr. Biol. 31, 192-197.e4 (2021).

21. Buzsáki, G. Hippocampal sharp wave-ripple: A cognitive biomarker for episodic memory and planning. Hippocampus 25, 1073–1188 (2015).

22. Nitzan, N., Swanson, R., Schmitz, D. & Buzsáki, G. Brain-wide interactions during hippocampal sharp wave ripples. Proc. Natl. Acad. Sci. 119, e2200931119 (2022).

23. O’Callaghan, C., Walpola, I. C. & Shine, J. M. Neuromodulation of the mind-wandering brain state: the interaction between neuromodulatory tone, sharp wave-ripples and spontaneous thought. Philos. Trans. R. Soc. B Biol. Sci. 376, 20190699 (2020).

24. International Brain Laboratory et al. Spike sorting pipeline for the International Brain Laboratory. Online resource at 10.6084/m9.figshare.19705522.v3 (2022).

25. Celada, P., Puig, M. V. & Artigas, F. Serotonin modulation of cortical neurons and networks. Front. Integr. Neurosci. 7, (2013).

26. Busse, L. et al. The Detection of Visual Contrast in the Behaving Mouse. J. Neurosci. 31, 11351–11361 (2011).

27. International Brain Laboratory et al. Standardized and reproducible measurement of decision-making in mice. eLife 10, e63711 (2021).

28. International Brain Laboratory et al. A Brain-Wide Map of Neural Activity during Complex Behaviour. 2023.07.04.547681 Preprint at 10.1101/2023.07.04.547681 (2023).

29. Fonseca, M. S., Murakami, M. & Mainen, Z. F. Activation of Dorsal Raphe Serotonergic Neurons Promotes Waiting but Is Not Reinforcing. Curr. Biol. 25, 306–315 (2015).

30. Miyazaki, K. W., Miyazaki, K. & Doya, K. Activation of Dorsal Raphe Serotonin Neurons Is Necessary for Waiting for Delayed Rewards. J. Neurosci. 32, 10451–10457 (2012).

31. Miyazaki, K., Miyazaki, K. W. & Doya, K. Activation of Dorsal Raphe Serotonin Neurons Underlies Waiting for Delayed Rewards. J. Neurosci. 31, 469–479 (2011).

32. Stemmler, M. & Koch, C. How voltage-dependent conductances can adapt to maximize the information encoded by neuronal firing rate. Nat. Neurosci. 2, 521–527 (1999).

33. Correia, P. A., Matias, S. & Mainen, Z. F. Stereotaxic Adeno-associated Virus Injection and Cannula Implantation in the Dorsal Raphe Nucleus of Mice. Bio-Protoc. 7, e2549 (2017).

34. Jackson, N. & Muthuswamy, J. Artificial dural sealant that allows multiple penetrations of implantable brain probes. J. Neurosci. Methods 171, 147–152 (2008).

35. Newman, J. P. et al. Optogenetic feedback control of neural activity. eLife 4, e07192 (2015).

36. International Brain Laboratory. IBL protocol for mice training. Preprint at 10.6084/m9.figshare.11634729.v5 (2022).

37. International Brain Laboratory et al. Standardized and reproducible measurement of decision-making in mice. eLife 10, e63711 (2021).

38. Jun, J. J. et al. Fully integrated silicon probes for high-density recording of neural activity. Nature 551, 232–236 (2017).

39. International Brain Laboratory. IBL protocol for labeling the tip of Neuropixels probes. Online resource at 10.6084/m9.figshare.19698130.v2 (2022).

40. International Brain Laboratory. IBL protocol for electrophysiology recording using Neuropixels probe. Online resource at 10.6084/m9.figshare.19697896.v2 (2022).

41. International Brain Laboratory. IBL protocol for perfusion and shipment of brain sample. Online resource at 10.6084/m9.figshare.19698061.v2 (2022).

42. International Brain Laboratory. IBL protocol for mouse brain reconstruction and registration. Online resource at 10.6084/m9.figshare.19698895.v2 (2022).

43. International Brain Laboratory. IBL protocol for registering the electrode location using LASAGNA. Online resource at 10.6084/m9.figshare.19698166.v2 (2022).

44. Mathis, A. et al. DeepLabCut: markerless pose estimation of user-defined body parts with deep learning. Nat. Neurosci. 21, 1281–1289 (2018).

45. Birman, D. et al. Video hardware and software for the International Brain Laboratory. Online resource at 10.6084/m9.figshare.19694452.v1 (2022).

46. Montijn, J. S. et al. A parameter-free statistical test for neuronal responsiveness. eLife 10, e71969 (2021).

47. Montijn, J. S., Meijer, G. T. & Heimel, J. A. A new family of statistical tests for neuronal spiking and autocorrelated timeseries data. 2023.10.30.564780 Preprint at 10.1101/2023.10.30.564780 (2024).

48. Seabold, S. & Perktold, J. Statsmodels: Econometric and Statistical Modeling with Python. Proc. 9th Python Sci. Conf. (2010) doi:10.25080/Majora-92bf1922-011.

49. Harris, K. D. Nonsense correlations in neuroscience. bioRxiv (2020) doi:10.1101/2020.11.29.402719.

