## Supplementary Figures for "Serotonin drives choice-independent reconfiguration of distributed neural activity"

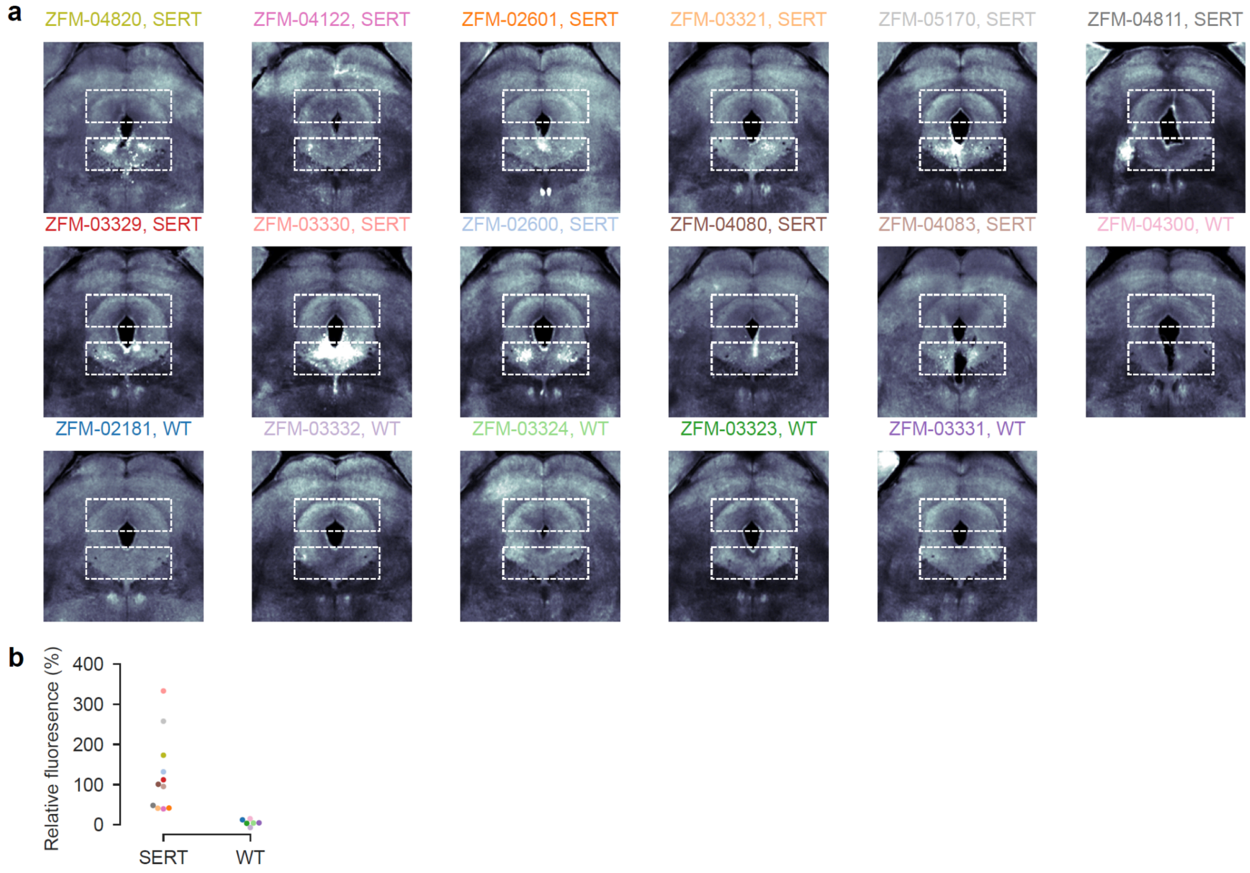


**Supplementary Figure 1. Channelrhodopsin expression is restricted to DRN of SERT-Cre mice. (a)** Coronal slices of the DRN of all mice in the study, imaged by a serial sectioning two-photon scanner. DRN fluorescence was defined as the sum of all the pixel values in the bottom dashed rectangle, an equal size control area was defined as the top rectangle. Relative fluorescence was calculated as the percentage fluorescence increase in DRN versus control area. The coronal slice with the highest relative fluorescence between -4.4 and -4.6 mm AP was chosen for analysis. Titles are mice names and whether the mouse was a SERT-cre or WT control. **(b)** Relative fluorescence, calculated as described in (a), for each mouse. The color of the dot corresponds to the color of the title in (a) to indicate from which mouse the value was from.

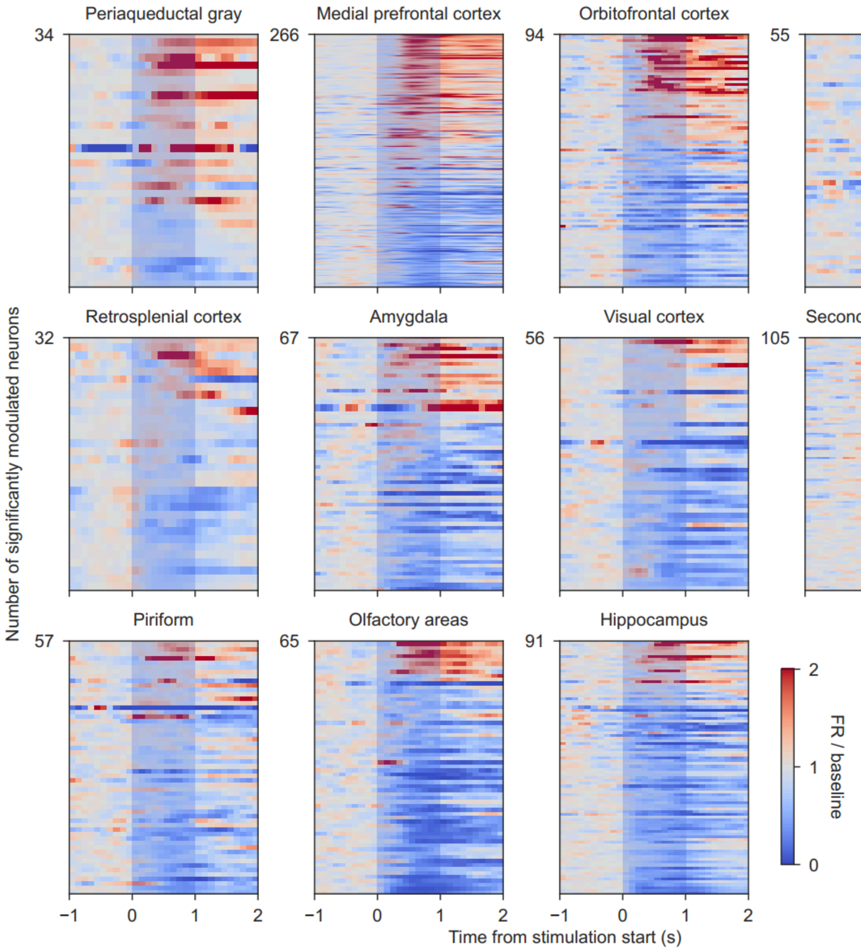


**Supplementary Figure 2. All significantly 5-HT modulated neurons during quiet wakefulness.** Each neuron is plotted as a colored row per brain region. The color indicates the mean firing rate over trials (as calculated in Fig. 2e) which is divided over the baseline (mean firing rate [-1 to 0s] + 0.1 spks/s) for plotting purposes. Light blue vertical bar between 0 and 1s indicates when the 5-HT stimulation was on.

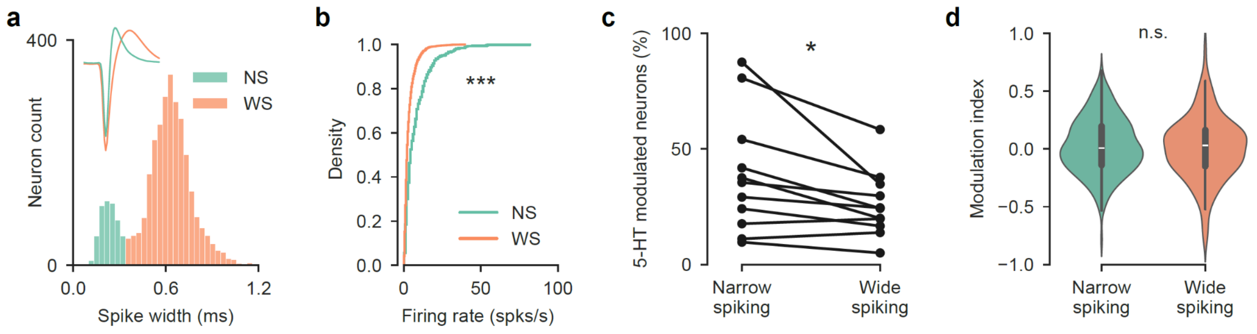


**Supplementary Figure 3. Waveform clustering separates regular spiking neurons from putative fast spiking interneurons. (a)** Spike width was used to classify a neuron as narrow (NS) or wide spiking (WS) by applying a cut-off value of 0.35 ms. The mean waveforms of the two groups of neurons are displayed in the inset. **(b)** The firing rate of narrow spiking neurons was significantly higher compared to wide spiking neurons, indicating narrow spiking neurons are putative fast-spiking interneurons. $p$ < 0.001, Kolomogorov-Smirnov test. **(c)** Slightly more narrow spiking (putative interneurons) than wide spiking neurons were significantly modulated by 5-HT stimulation. Each line is an individual mouse. $p$ = 0.02, paired t-test. **(d)** The modulation index of narrow and wide spiking neurons was not significantly different. Plotted are violin plots of the distributions of modulation indices for all narrow and wide spiking neurons. $p$ = 0.43, independent samples t-test.

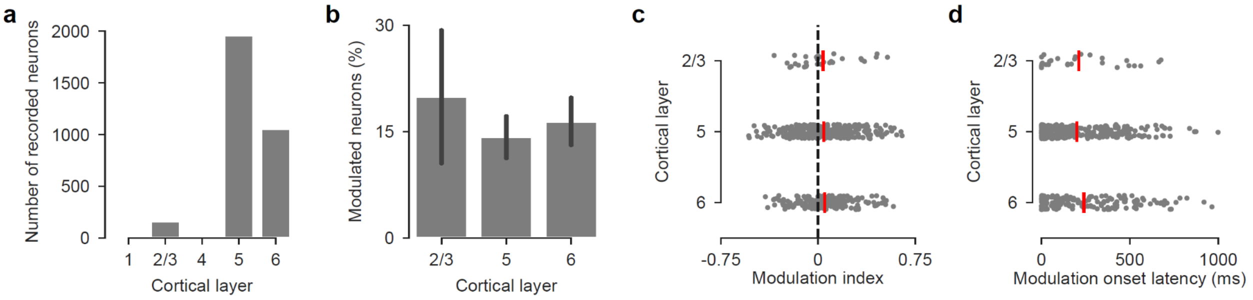


**Supplementary Figure 4. Serotonergic modulation does not show layer dependent differences. (a)** The total number of recorded neurons per cortical layer across all cortical recordings. **(b)** The percentage of modulated neurons per cortical layer. Bars show mean over animals and error bars the standard error of the mean. **(c)** The modulation index of each recorded neuron (gray dots) with the mean over neurons shown as a red vertical stripe. **(d)** The modulation onset latency as in (c) per cortical layer.

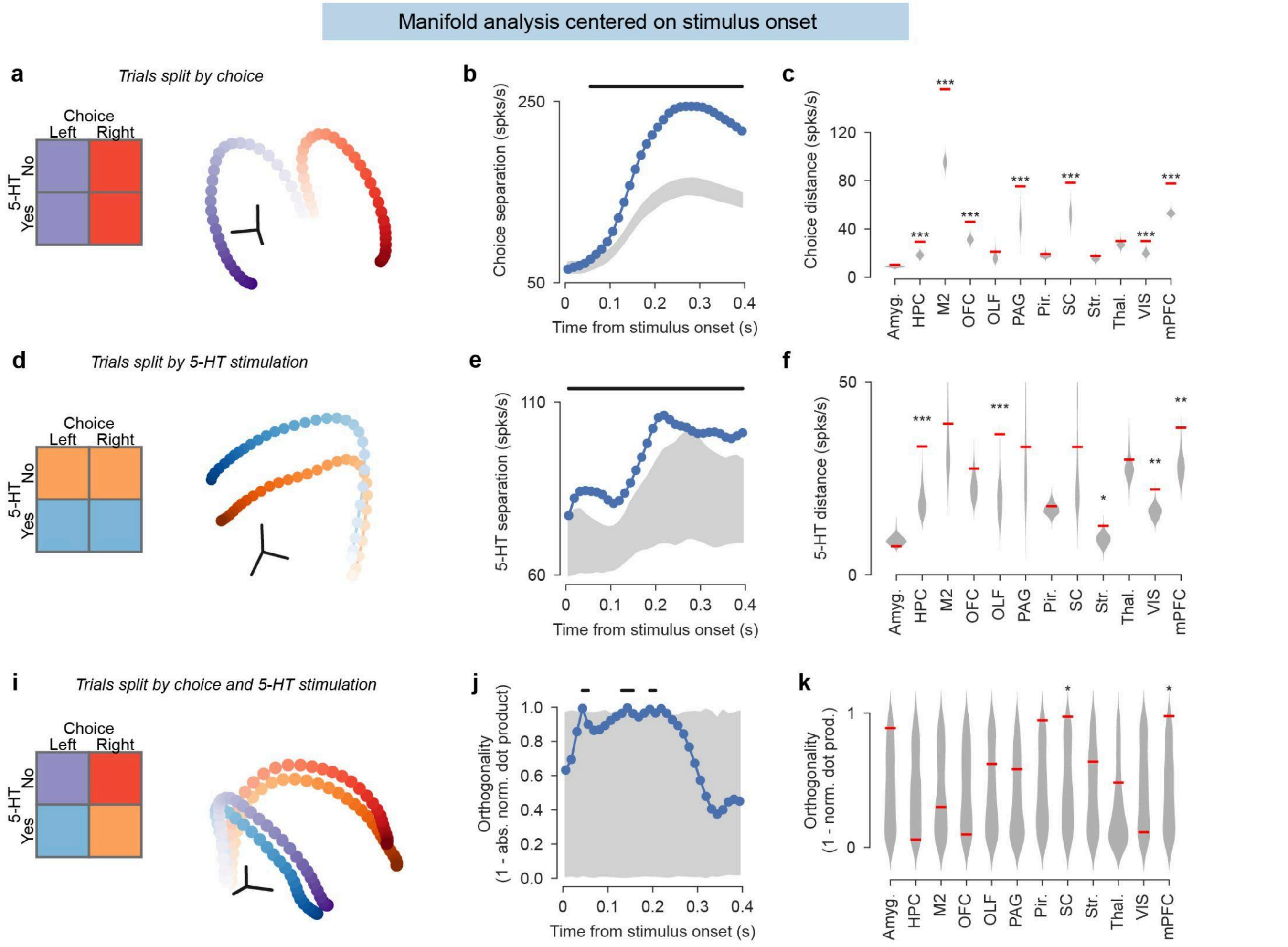


**Supplementary Figure 5. Manifold analysis centered on stimulus onset. (a,b,d,e,i,j)** Panels as in Figure 5 but centered on stimulus onset instead of choice point.
